# CRISPR-Stabilized Voronoi-like Domains in Synthetic Bacterial Populations

**DOI:** 10.1101/2024.07.31.606050

**Authors:** Daniela Moreno, Yerko Ortiz, Diego Araya, Gianna Arencibia, Fivos Panetsos, Martín Gutiérrez

## Abstract

Synthetic multicellular systems provide a controllable framework for studying how local interactions generate robust spatial organization in growing populations. Pattern formation is approached here as a programmable design problem, in which signaling, growth, and commitment must be coordinated to produce stable collective structure rather than treated as a secondary outcome. Voronoi-like tilings represent a useful class of organizations in which domains remain locally exclusive, collectively cover space, and often converge toward efficient hexagonal geometries, yet they have rarely been examined as emergent collective states governed by explicit local rules and tunable spatial control parameters.

Here, a spatial-resolution-based framework is presented to generate stable Voronoi-like partitions in simulated bacterial colonies by combining quorum sensing, leader-follower role assignment, band-detection logic, and CRISPR-Cas9-mediated irreversible commitment. The system assigns local node identities, stabilizes spatial domains over time, and produces Voronoi-like partitions that emerge from distributed interactions rather than centralized control. The framework is implemented in the gro agent-based simulator and analyzed across parameter sweeps that vary leader-signal thresholds, node-size control, and signal propagation features. Robust spatial organization is found to depend on irreversible commitment, appropriate threshold separation, and controlled diffusion and degradation, revealing distinct robustness regimes that separate clean Voronoi-like partitions from deformed or mixed-identity patterns. These results position Voronoi-like spatial partitioning as a representative problem for programmable microbial active matter and support the use of synthetic gene circuits as minimal platforms for studying and exploiting spatial self-organization in microbial populations.

## 1. Introduction

Synthetic multicellular systems provide a controllable framework for studying how local interactions generate robust spatial organization in growing populations. In this context, pattern formation is not only a developmental phenomenon but also a programmable design problem in which signaling, growth, and commitment must be coordinated to produce stable collective structure.

Most synthetic patterning studies have focused on reaction-diffusion mechanisms, morphogen gradients, lateral inhibition, or collective motility. By contrast, much less attention has been given to Voronoi-like partitioning as an emergent collective state governed by explicit local rules and tunable spatial control parameters. This question is relevant because Voronoi-like tilings capture a useful class of spatial organizations in which domains remain locally exclusive, collectively cover space, and often converge toward efficient geometries such as hexagonal arrangements. In microbial systems, achieving such organization requires more than signal exchange alone; it also requires mechanisms that define node identity, prevent unstable reassignment, and maintain separation between equivalent neighboring states.

The present work addresses this problem through a minimal agent-based framework for programmable Voronoi-like partitioning in bacterial colonies. The proposed architecture combines quorum sensing, leader-follower assignment, band-detection logic, and CRISPR-Cas9-mediated irreversible commitment to transform local communication into stable spatial domains.

The originality of the study lies in two aspects. First, it reframes Voronoi-like pattern formation as a minimal active-matter problem in synthetic biology, asking which local rules and control parameters are sufficient to generate robust spatial partitions^20,23-25^. Second, it proposes a concrete modular genetic architecture in which irreversible commitment stabilizes node identities and reveals an explicit trade-off between robustness and plasticity. Beyond generating visually recognizable patterns, the framework enables quantitative analysis of spacing, clustering, and geometric robustness across parameter regimes. This positions the system not only as a synthetic design strategy for spatial task allocation in microbial populations, but also as a comparative model for studying general principles of biological self-organization and bioinspired partitioning.

In our study we identify minimal design rules for robust Voronoi-like spatial partitioning in a synthetic multicellular system. First, a spatial-resolution-based framework is introduced that uses quorum sensing, leader–follower dynamics, and irreversible genetic commitment to generate stable Voronoi-like partitions in an agent-based bacterial colony model. Second, the study analyzes how leader-signal thresholding, node-size control, and commitment logic jointly influence domain stability, spacing, and emergent geometry^9,14^. By framing the problem in terms of node identity, robustness regimes, and explicit spatial control parameters rather than exact geometric construction, this work connects synthetic pattern formation with a broader agenda of programmable collective organization in microbial active matter^6,16,36^.

## 2. Background and state of the art

Developmental biology is a pluripotent field that relates to disciplines such as genetics and immunology while preserving its own identity^1^. In microbiology, developmental biology can be understood as the capacity for self-organization and self-regulation in a biological system^2^. The term pattern refers to coordinated differential gene expression over time and space, where such patterns lead to variations in cell behavior that drive morphogenesis^3^. Studying self-organization is therefore essential for understanding the evolution of biocomplexity^4^. A central example is embryonic development, in which self-organizing mechanisms drive the transition from undifferentiated cells to a structured organism through pattern establishment, cell-state determination, and the emergence of physical form^5^. These processes depend on genetic regulation, changes in cell shape, cell-cell interaction, proliferation, growth, and movement^3,6,7^.

Pattern generation depends on multiple factors. Cell morphology can strongly influence cell lineage fate^8^, and pattern formation can also be modulated by protein degradation and repressor expression ratios^9^. Previous studies have described several mechanisms for biological pattern generation, including symmetry breaking^4^, reaction-diffusion dynamics^10,11^, morphogen gradients^12^, organized bacterial movement^13^, cell-cell communication^14^, and nutrient heterogeneity^15^. These examples show that spatial organization can emerge from local interactions, long-range signaling, or combinations of both.

In synthetic developmental biology, developmental processes are combined with synthetic biology to engineer and control multicellular behavior, thereby improving both mechanistic understanding and functional design^16^. Synthetic systems provide simplified and controllable settings in which multicellular coordination can be studied with fewer confounding factors^6^. This is especially relevant because one of the central challenges in multicellular systems is achieving coordinated behavior under increasing complexity^17^. Synthetic biology addresses this challenge by enabling cells to execute predefined functions through programmable genetic modules^17,18^. Constructed multicellular systems also provide a basis for advances in tissue engineering, biomaterials, and biosensing^19^, while individual-based models help analyze emergent properties of genetic circuits in multicellular environments^9^.

Alternative approaches based on DNA computing and protein filament networks have also been proposed to address complex collective problems^4,13,14^. These biologically based solutions exploit massive parallelism and local interactions to implement nontrivial computations. However, existing strategies often provide limited control over spatial resolution and over the explicit local rules that determine how stable spatial domains emerge, persist, and reorganize in growing cellular populations^5,18^. In many cases, spatial structure is treated as a secondary outcome of molecular computation rather than as a primary design target.

This limitation becomes especially important when spatial organization is interpreted not only as a developmental or computational outcome but also as a problem of active collective matter under programmable constraints. In microbial populations, spatial domains arise from local sensing, diffusive communication, growth, and commitment processes acting without centralized coordination^2,6,14^. Under this view, the relevant question is not only whether a target pattern can be generated, but which minimal local interactions and control parameters determine the robustness, stability, and geometry of the resulting collective state. Recent work has shown that collective behavior in chemotactic microbial populations can be understood within a minimal agent-based framework in which robustness depends on explicit environmental control parameters rather than on signal amplitude alone^36^. A similar shift in emphasis can be applied to spatial partitioning problems.

The same perspective is adopted here. Rather than focusing on the construction of an exact Voronoi diagram, this work treats Voronoi-like spatial partitioning as a model problem for programmable multicellular organization. The proposed system assigns local node identities to bacterial groups, uses leader-follower logic and band detection to define and subdivide domains, and employs CRISPR-Cas9-based genetic commitment to stabilize assignments over time^20,23^. As a result, the colony self-organizes into stable domains separated by interfaces that approximate a Voronoi-like partition, and the quality of the pattern can be related to parameters such as signal thresholds, interaction ranges, and decision timing^9,21^.

## 3. System description

The system implements a distributed strategy for generating a Voronoi-like partition in a bacterial colony. Instead of treating the problem purely as graph coloring, the framework assigns a node identity to groups of bacteria that behave as local spatial domains. Adjacent domains must maintain distinct identities, and these identities are established and preserved through intercellular communication and irreversible genetic commitment. The system builds on previously described intercellular metaheuristic frameworks and uses the Moore neighborhood concept to define local interaction ranges through diffusive autoinducers, fission, and signal degradation parameters^21^. In this formulation, nodes are interpreted as spatially bounded subpopulations with coherent identity and behavior, and the Moore-neighborhood interaction range becomes an explicit control parameter for how local signaling shapes the resulting partition^21^.

### 3.1. Leader-follower dynamic

Although the Moore neighborhood specifies the interaction range between neighboring subpopulations, it does not explicitly define node size in the colony model. To establish clear node boundaries, a fixed reference signal, termed the leader signal, is introduced. Its intensity encodes distance from a designated leader bacterium, propagates through the colony, and defines the spatial extent of each node independently of neighborhood-based signals^21^ (see Figure 1). Node organization follows a leader-follower dynamic in which leaders act as local organizers by emitting a signal associated with a specific node identity. Surrounding bacteria detect this signal, adopt the leader’s node identity, and become followers of that domain. To prevent reassignment, followers eliminate plasmids that would otherwise allow them to respond to leaders of alternative identities, thereby ensuring exclusive and permanent association^23^.

**Figure 1.**
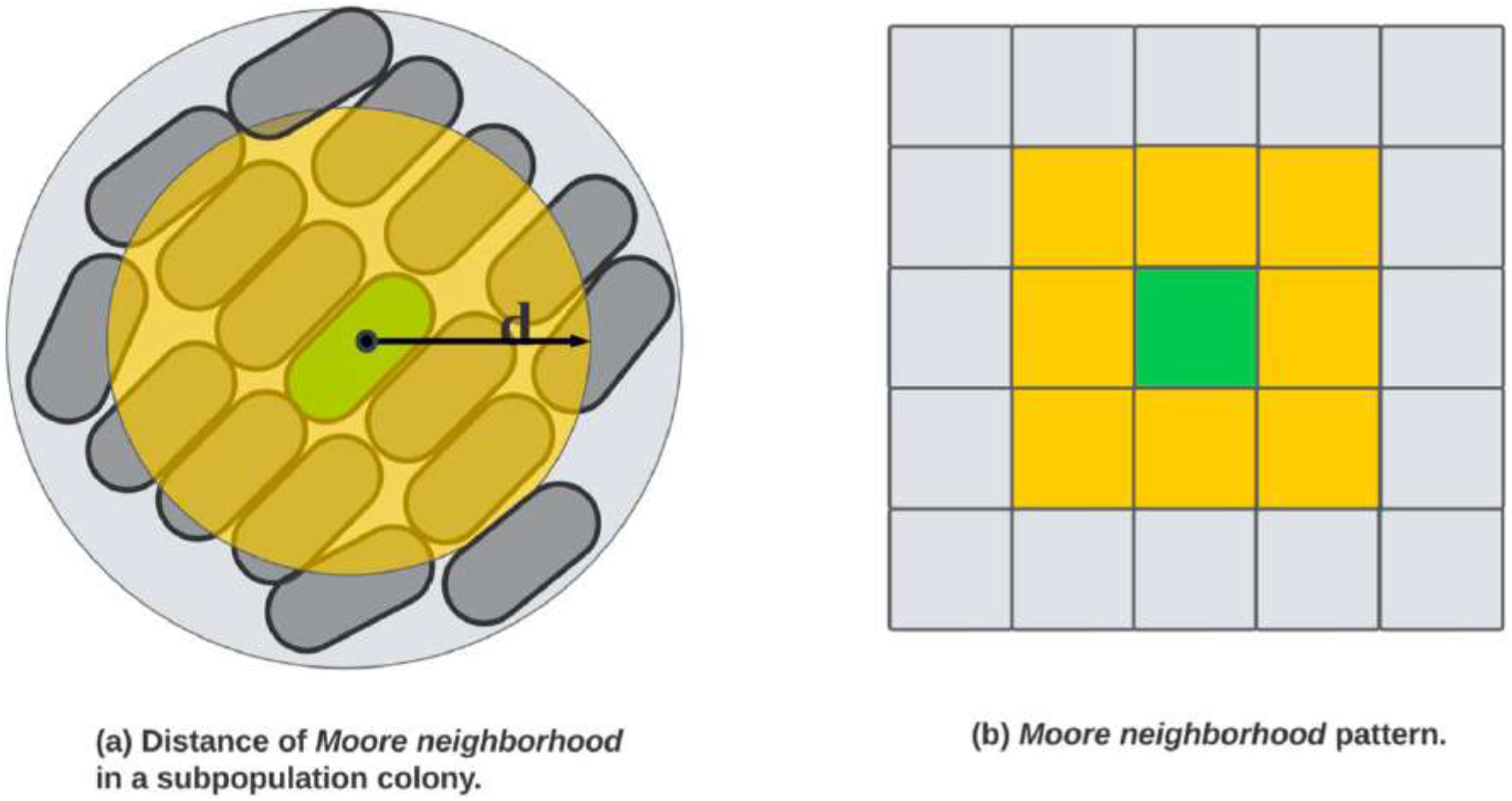
Node definition through neighborhood range and leader-centered signaling. Left, schematic representation of neighborhood range in a bacterial colony, showing how local interaction distance can be interpreted spatially in a multicellular setting. Right, theoretical Moore-neighborhood pattern used as inspiration for delimiting local interaction range. In the proposed system, this neighborhood concept is implemented biologically through leader-emitted quorum-sensing signals, whose diffusion, degradation, and detection thresholds define the effective spatial extent of a node rather than a purely geometric adjacency rule.

Interactions between neighboring domains are coordinated through a local identity-exclusion logic in which adjacent nodes must not share the same identity. Leaders select identities using greedy local rules based on neighboring information, which yields a Voronoi-like partition of the colony. Once a bacterium commits to a leader, its node membership remains fixed, stabilizing domain boundaries and preserving the partition over time^21,23^.

### 3.2. Expected behavior

The interaction between predefined roles leads to a sequence of stages that collectively generate the colony-scale pattern: signal initiation, node definition, leader-signal intersection, node-identity assignment, and iterative repetition until the colony becomes fully partitioned^21^.

At signal initiation, two leader bacteria are introduced into the colony, each emitting a unique signal that establishes the preliminary boundaries of its node. During node definition, leaders continue broadcasting their signals, thereby limiting the size of their respective nodes, while bacteria that detect a leader signal activate a genetic circuit that prevents them from becoming new leaders and commits them to that node identity. As leader signals expand, overlaps occur and new boundaries emerge; different pairs of intersecting identities trigger the appearance of new leaders carrying complementary node identities. Each new leader establishes a new domain that affects neighboring regions through additional signal propagation and leader selection. These processes then repeat, causing nodes to subdivide and colony regions to continue acquiring identities until a cohesive Voronoi-like partition is formed^21^. The whole process is shown in Figure 2.

**Figure 2.**
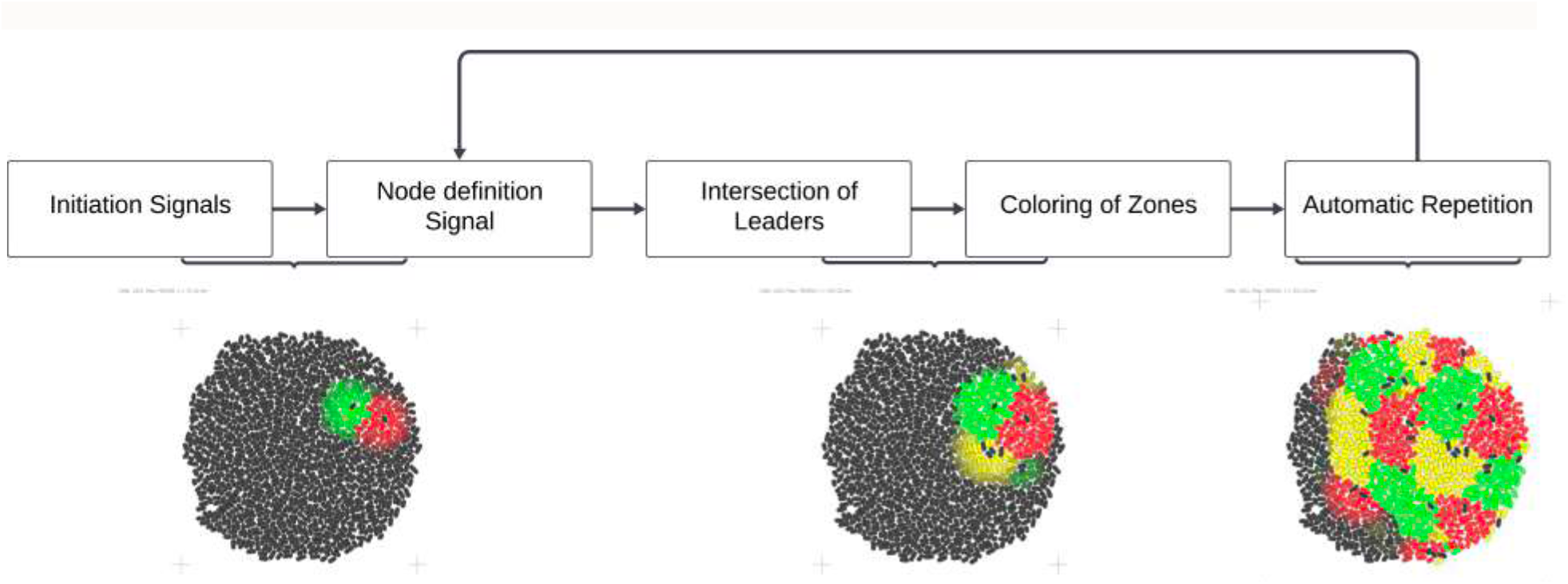
Expected collective behavior of the spatial partitioning program. Schematic sequence of the expected colony-level dynamics: signal initiation by the first leaders, node definition through leader-derived quorum-sensing fields, leader emergence at signal-intersection zones, assignment of node identity to surrounding followers, and recursive repetition of the process. Colors indicate node identity labels only; the underlying spatial organization is generated by diffusible signaling and threshold-based response.

### 3.3. Circuit design and implementation

The system is implemented as a genetic circuit that enables distributed computation within a bacterial colony. The design is defined at an abstract level using generic identifiers for signals, promoters, plasmids, genes, and proteins, which facilitates reasoning about circuit functionality independently of specific biological components and enables later implementation with automated design tools such as CELLO^22^. To address node-size limitation and preserve stable assignments, the design uses CRISPR-Cas9 within a modular plasmid-based architecture^23^. The circuit is divided into functionally grouped plasmids that can be selectively deleted to prevent the formation of additional leaders in follower cells while preserving the remaining circuit functions. The same global architecture is replicated for each node identity, differing only in identity-specific modules.

The plasmid modules are organized as follows. pLeader selects leader bacteria and eliminates follower-active modules related to identity expression and leader-signal repression, thereby stabilizing leader identity and enabling emission of the leader signal that defines the local follower zone. pRepLeaderX represses leader behavior in follower cells, preventing them from adopting a leader role and allowing expression of the node-identity program. pRepColorX suppresses the “X” identity program when two other “X” node zones are in close proximity through band-detection logic, acting as an exclusion rule between nearby equivalent identities. pColorX operates in the leader bacterium and, via quorum sensing, induces the corresponding node-identity program in follower cells. pBand, activated by the leader signal, defines the band that bounds the node zone and neighboring regions. pInit implements the band-detection mechanism; the overlap of adjacent zones carrying two different node identities activates this plasmid and designates a bacterium as the new leader for a newly formed node. Finally, pDead ensures irreversible commitment: upon reception of the leader signal, Cas9 deletes pLeader, pDead, and identity-specific leader modules in follower cells, preventing the later emergence of additional leaders^20,23^. A complete diagram of the circuit is shown in Figure 3.

**Figure 3.**
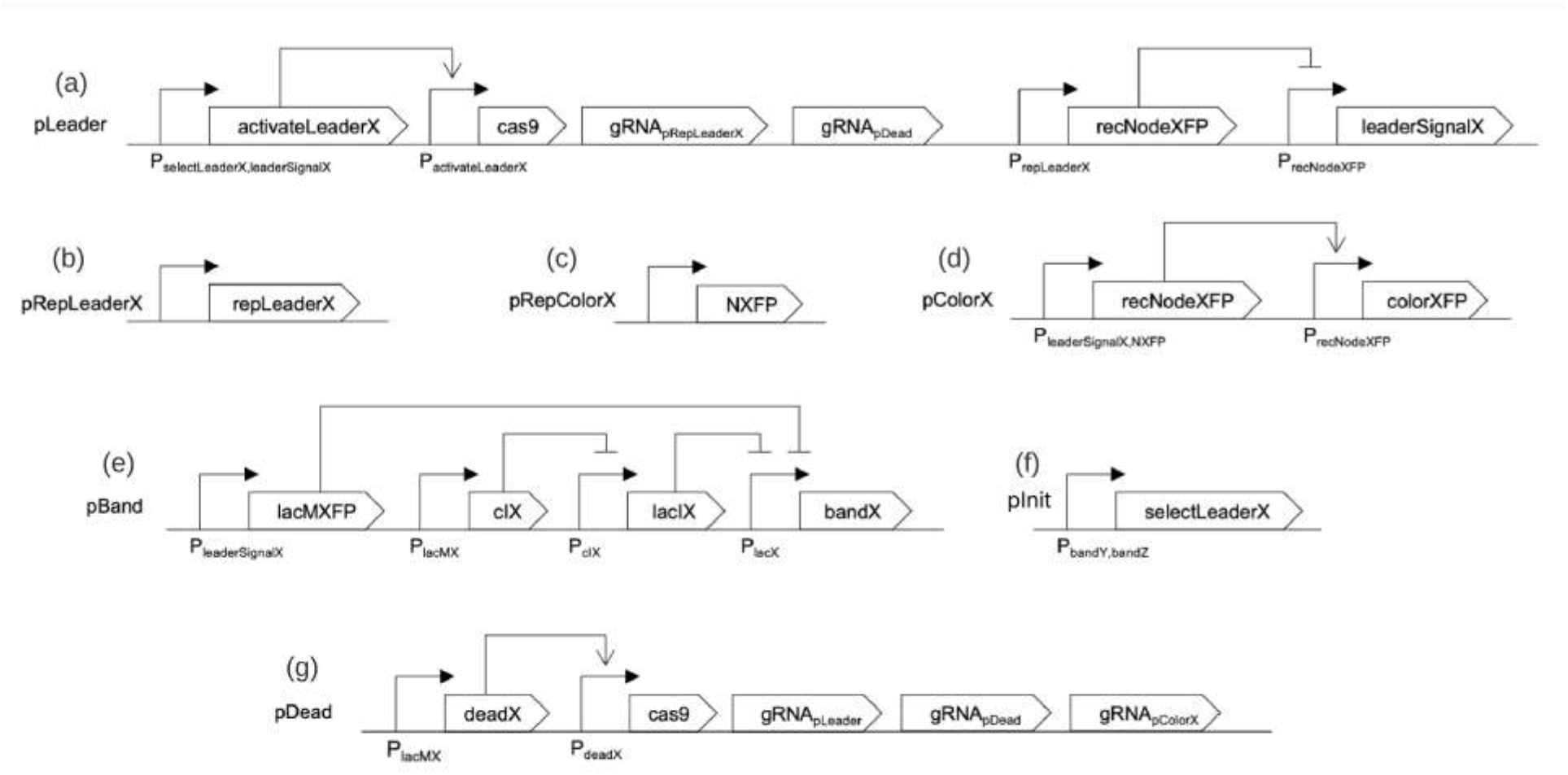
Modular genetic architecture for leader formation, node labeling, and irreversible commitment. Global architecture of the genetic circuit used to implement distributed spatial partitioning. The same circuit logic is repeated for each node identity, with X denoting the focal node label and Y and Z the alternative labels. Modules control leader selection, repression of competing leader states, node-identity readout, band detection, recursive leader initiation at signal-intersection zones, and CRISPR-Cas9-mediated plasmid deletion for irreversible commitment. Together, these modules transform transient quorum-sensing exposure into stable node identity and persistent spatial organization.

### 3.4. Convergence to the solution

The evaluation function is based on convergence through intersections between neighboring node identities, following a greedy local resolution procedure. When a bacterium becomes part of a node, it converges to the corresponding node identity within that group. Once all bacteria in the colony have adopted a node identity, excluding the initiating leaders, the colony is considered to have reached a stable spatial partition.

The quality of the solution is not evaluated through regression or supervised training, but through the collective outcome of local interactions. Each bacterium performs its designated function according to its local environment and contributes, through quorum sensing, to a coherent colony-scale organization. Within this framework, the observed pattern is interpreted as one admissible global partition selected by the local greedy rules, providing a concrete instance of how distributed decision making and explicit control parameters can jointly determine a Voronoi-like spatial organization without centralized optimization^20,21^.

### 3.5. gro simulation

To evaluate the proposed framework, simulations were implemented using gro, an agent-based simulator for bacterial behavior that supports CRISPR-Cas9 logic and quorum-sensing signaling^24,25^. The simulation is based on a band-detection genetic circuit operating with two signal-sensitivity thresholds to detect sufficient spatial separation between nodes while implementing the leader-follower designation mechanism^19,26^. The initial population consisted of one red initiator, one green initiator, and 500 bacteria carrying plasmids that enable them to become leaders or followers. Under this configuration, the desired spatial pattern emerges after appropriate parameter tuning.

In addition to visual outputs, the simulation generates a CSV file containing the spatial position and node-identity assignment of each bacterium. These data were used for quantitative spatial analysis, including density-based clustering and centroid-distance measurements.

## 4. Results

The results from gro simulations are presented in terms of node-identity persistence, node-size sensitivity, and spatial organization, including a quantitative analysis of same-identity spacing across the colony.

### 4.1. Node-identity persistence

Nodes contain a limited number of bacteria within each spatial zone. Once a bacterium commits to a node identity, it does not switch to an alternative identity, resulting in stable assignment across the colony. In the static configuration (Figure 4(a)), the colony exhibits well-defined spatial patches, with each region assigned one of the available node identities and adjacent nodes maintaining distinct identities, producing clearly separated zones with negligible overlap.

**Figure 4.**
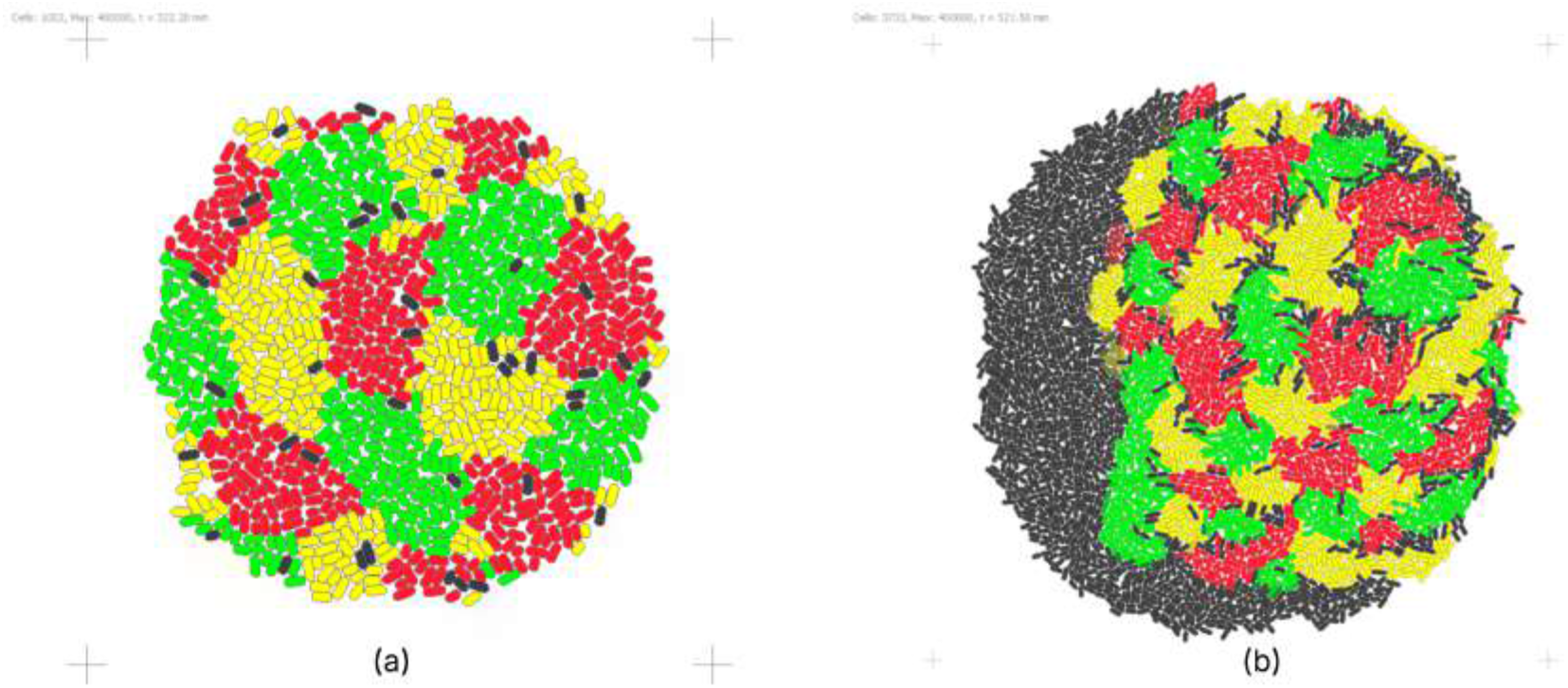
Node-identity persistence and preservation of spatial partitioning under static and growing colony conditions. Representative GRO simulations showing the stability of node identity after local commitment. (a) In the static colony, the system generates well-separated spatial domains with persistent node labels and clear interfaces between adjacent regions. (b) When colony growth is included, the global partition remains preserved despite local deformations and gaps caused by expansion. In both cases, the results indicate that CRISPR-mediated irreversible commitment stabilizes follower identity after quorum-sensing-based assignment, preventing reassignment even as colony geometry changes.

When bacterial growth is incorporated (Figure 4(b)), the global spatial organization is preserved: colony expansion introduces gaps and local deformations in patch shape, but node assignment and overall identity distribution remain consistent. This behavior indicates that the partition is stabilized by the genetic commitment mechanism and is robust to growth-induced perturbations^23-25^.

### 4.2. Node-size parameter sensitivity

A parameter sweep was conducted to examine the effect of leader-signal sensitivity detection on node size. The low sensitivity threshold was kept fixed, whereas the high sensitivity threshold was varied, thereby changing the size of the death zone and the effective spatial extent of each node. This produced clear differences in domain size and geometry.

These results show that node size is an emergent property controlled by threshold-dependent spatial decoding of the leader signal. Larger domains improve coverage but are more prone to deformation due to interactions with neighboring nodes, whereas smaller domains better preserve local regularity (see Figure 5). Threshold spacing therefore emerges as a key control parameter governing the trade-off between coverage and geometric robustness^9,14^.

**Figure 5.**
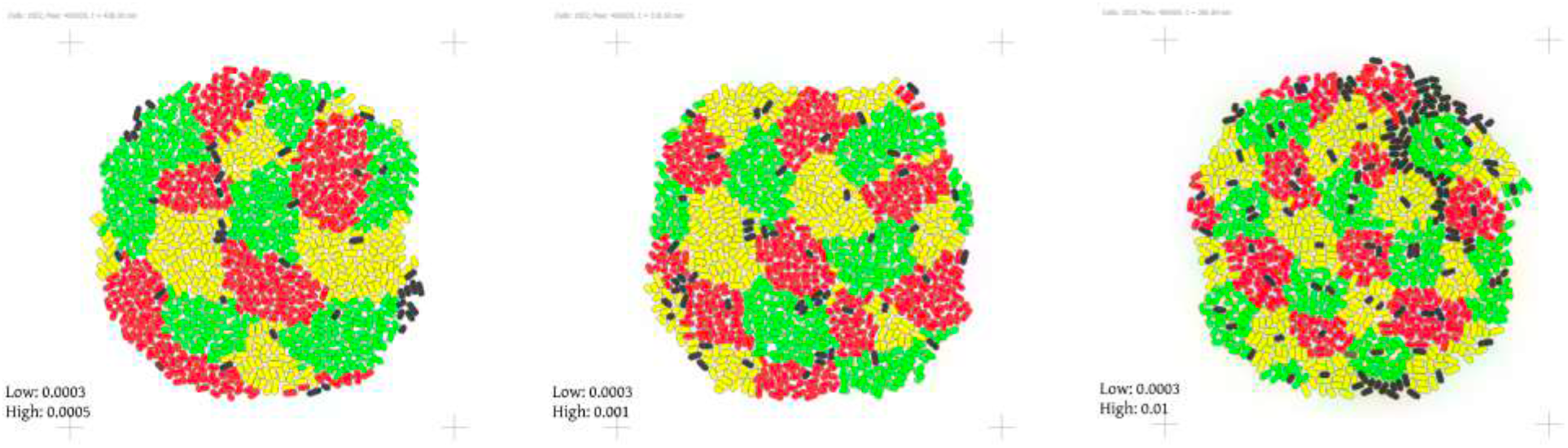
Sensitivity of node size to leader-signal detection thresholds. Parameter sweep showing the effect of varying the high leader-signal detection threshold while keeping the low threshold fixed. This manipulation changes the width of the effective response band and therefore modifies the spatial extent of node domains. Larger threshold separations produce broader domains but also increase deformation due to node collisions, whereas narrower bands constrain node growth and preserve more regular local geometry. The figure illustrates that node size is an emergent property controlled by quorum-sensing signal interpretation rather than by an externally imposed geometric radius.

### 4.3. Spatial clustering analysis of node organization

To quantify the spatial arrangement of committed cells in the final colony state, exported gro coordinates were analyzed using DBSCAN after grouping bacteria by node-identity label. This allowed coherent domains to be identified independently for each identity class and provided a compact geometric description of the emergent partition via cluster centroids and inter-centroid distances. Because the analysis was performed separately for each identity, it specifically tested whether nodes sharing the same label remained spatially separated rather than merging into contiguous regions.

The clustering results indicate that each node-identity class forms multiple distinct spatial domains across the colony. GFP-expressing cells formed 5 clusters with a minimum centroid-to-centroid distance of 16.78 µm, RFP-expressing cells formed 7 clusters with a minimum centroid distance of 15.46 µm, and YFP-expressing cells formed 10 clusters with a minimum centroid distance of 12.83 µm. These distances show that domains sharing the same identity are not adjacent but remain separated by a measurable spatial interval. The analysis is shown in Figure 6.

**Figure 6.**
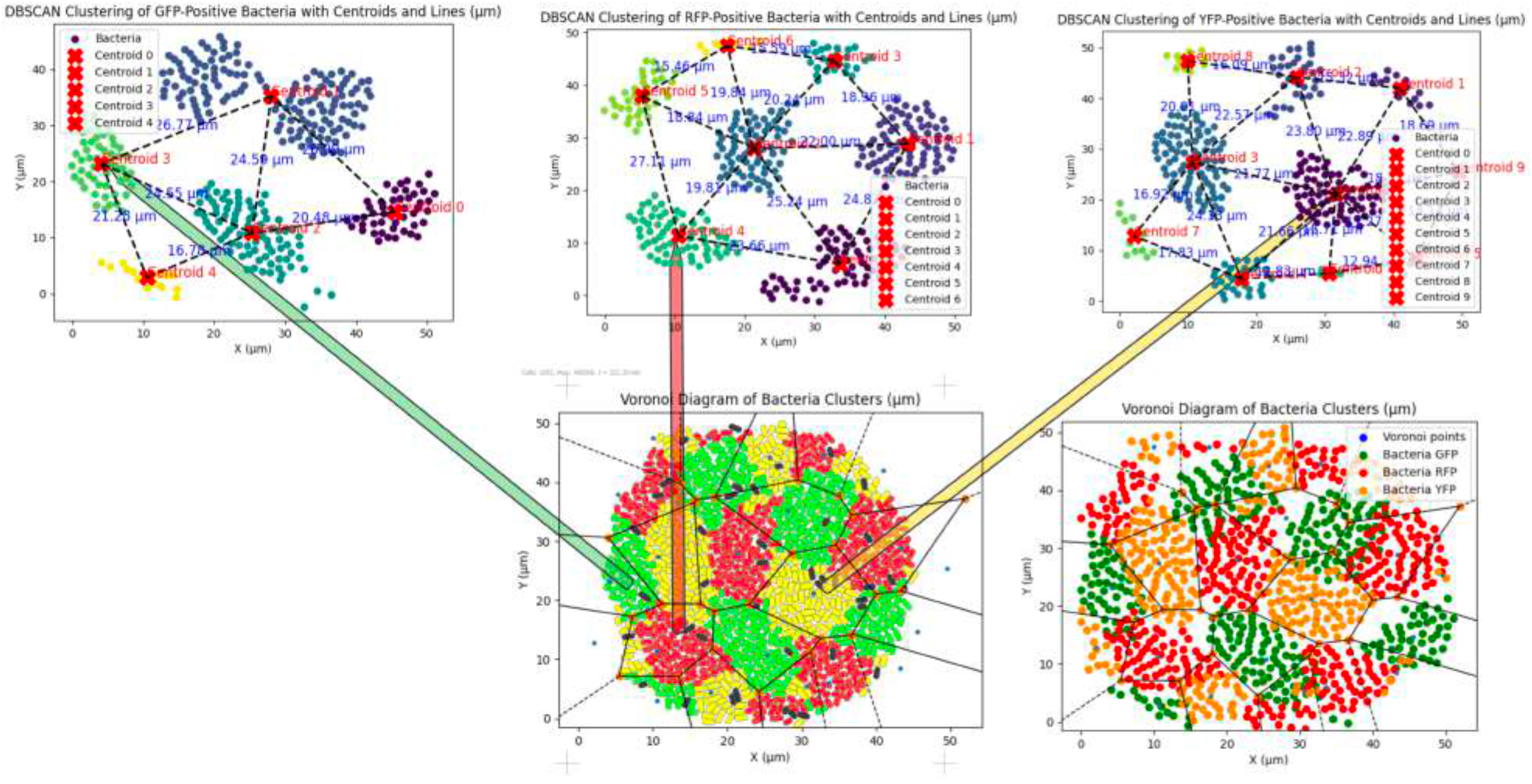
Spatial clustering analysis reveals separation between nodes sharing the same identity. DBSCAN-based clustering of bacterial positions extracted from GRO simulations, grouped by node-identity label. The analysis identifies dense clusters corresponding to spatially coherent node domains and estimates centroid-to-centroid distances among clusters of the same label. The observed minimum distances between like-labeled clusters indicate that the system maintains spatial separation among equivalent node identities, consistent with a local exclusion mechanism generated by quorum-sensing interactions and recursive leader formation.

This separation is consistent with the quorum-sensing and leader-follower logic of the system. Once a local domain is established, follower commitment and recursive leader formation at signal-intersection zones favor the emergence of neighboring domains with alternative identities, preventing immediate repetition of the same node state in adjacent regions. The observed spacing can thus be interpreted as an effective exclusion rule among equivalent node identities that is implemented implicitly by local signaling and commitment, rather than by any externally imposed geometric constraint^20,21^.

## 5. Discussion

This work identifies a minimal set of local mechanisms that are sufficient to generate robust Voronoi-like spatial partitioning in a growing bacterial colony. Rather than depending on centralized coordination, the final pattern emerges from the interaction between diffusible signaling, node-identity persistence, threshold-dependent size control, and decision timing, which together define the robustness regime of the collective spatial state^9,14,21^.

Signal dynamics played a central role in boundary formation. Proper calibration of diffusion, degradation, and threshold separation was required to produce clear node boundaries and adequate spatial resolution. Maintaining sufficient separation between low and high detection thresholds improved the sharpness of signal-defined borders and increased pattern clarity^9,14^.

Node-identity stabilization proved essential for preserving the partition over time. Without irreversible commitment, fluctuating signal levels would permit reassignment, generating instability and boundary erosion. CRISPR-Cas9-mediated plasmid deletion effectively prevented such reversals by locking each node into a single identity state, although this stabilization came at the cost of reduced adaptability once a decision had been made^23^.

Node size emerged as another important control parameter. Larger nodes improved territorial coverage but were more susceptible to deformation during collisions with neighboring domains, whereas smaller nodes better preserved local regularity. This trade-off indicates that signal-threshold tuning is central to balancing geometric stability against effective spatial coverage^9^.

Decision timing also influenced pattern quality. If neighboring nodes attempted to establish leadership simultaneously, conflicts and overlapping leadership events could reduce pattern clarity. Introducing temporal separation in activation and repressing the possibility of becoming a leader immediately after leader-signal emission, reduced these conflicts and improved exclusivity of local domains^21^.

Finally, the geometry of the resulting partition tended toward recurrent hexagonal organization. This is consistent with isotropic diffusion, approximately symmetric node spacing, and local exclusion between neighboring identities. The resulting geometry resembles efficient surface tilings observed in natural systems, supporting the view that the pattern arises from physically and biologically interpretable local constraints rather than from an externally imposed geometric template^27,33,34^ (see Figure 7).

**Figure 7.**
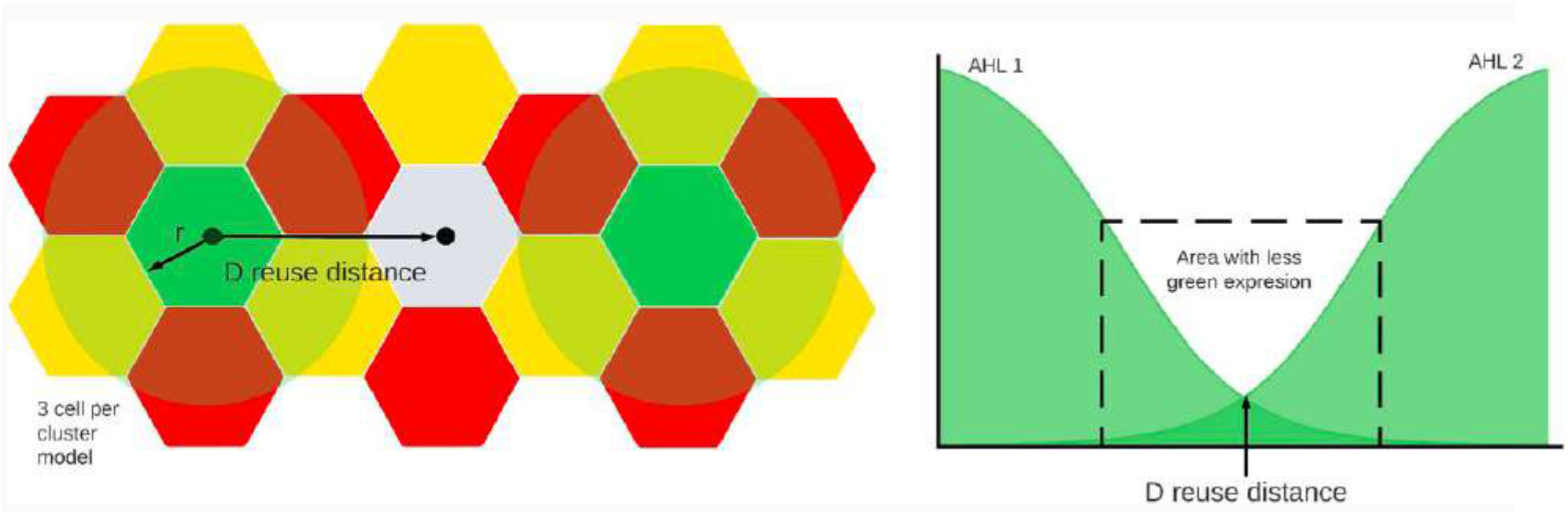
Gene expression decreasing with distance. In this configuration, three clusters of cells are shown, with green zones on the first and last clusters that emit quorum-sensing signals AHL1 and AHL2. The reuse distance defines the minimal separation at which the AHL signal associated with green coloring remains strong enough to reproducibly induce the same color while avoiding interference from background noise within the same signal^32^.

## 6. Applications

The ability to generate stable Voronoi-like partitions through local signaling and genetic commitment suggests applications wherever distributed control of spatial domains is required in multicellular systems. In this framework, Voronoi-like geometry is not only a convenient descriptor but a concrete target organization that can be approximated by realistic microbial systems under explicit control constraints, providing a testbed for design rules that link local genetic circuits to global tiling statistics^27-30^.

### 6.1 Spatial patterning in biological and synthetic systems

Voronoi-like structures appear in multiple biological contexts, such as epithelia where cell shapes and nuclear positions define tessellations that can be analyzed with Voronoi-based metrics^27^. The present framework offers a complementary synthetic approach in which a defined genetic circuit and quorum-sensing architecture generate analogous partitions in a controllable microbial setting, enabling systematic exploration of how local rules affect global tiling statistics^5,16,18^.

### 6.2 Programmable spatial task allocation in microbial collectives

Node identities can be coupled to distinct genetic functions rather than only to fluorescent reporters. Different domains within the colony could express complementary metabolic pathways, secrete distinct biomaterials, or implement separate sensing and actuation modules while maintaining stable spatial segregation through the same leader-follower and CRISPR-based commitment logic. The partition then defines a programmable map of functional territories in the population^6,16,19^.

### 6.3 Pattern-on-materials and biofabrication

The method has potential implications for biofabrication and pattern-on-materials applications, where spatially organized living systems are used to produce structures with defined properties. Voronoi-like partitions are associated with efficient packing and, in some cases, favorable mechanical organization; implementing such partitions through synthetic gene circuits could enable controlled deposition of polymers, pigments, or other products in efficiently packed arrangements^27,33^. The observed ability to reuse the same quorum-sensing signal across nonadjacent domains suggests scalable schemes in which a limited signal set supports multiple separated functional regions, an important consideration given the limited number of orthogonal quorum-sensing systems^14,35^.

### 6.4 Synthetic pattern formation as a comparative tool

Synthetic pattern-formation frameworks such as the one presented here can be used as comparative tools to probe which minimal local rules are sufficient to reproduce specific spatial statistics observed in natural systems. By varying commitment strength, threshold spacing, and decision timing within the same Voronoi-like architecture, it becomes possible to map how different parameter regimes recapitulate or deviate from epithelial tessellations, insect nest patterns, or other naturally occurring tilings, thereby linking abstract models of pattern formation to experimentally testable multicellular designs and explicitly characterizing robustness regimes in space^5,16,18,27,33,34^. The expected evolution of the geometry of the cell colonies to reach the mentioned patterns is shown in Figure 8.

**Figure 8.**
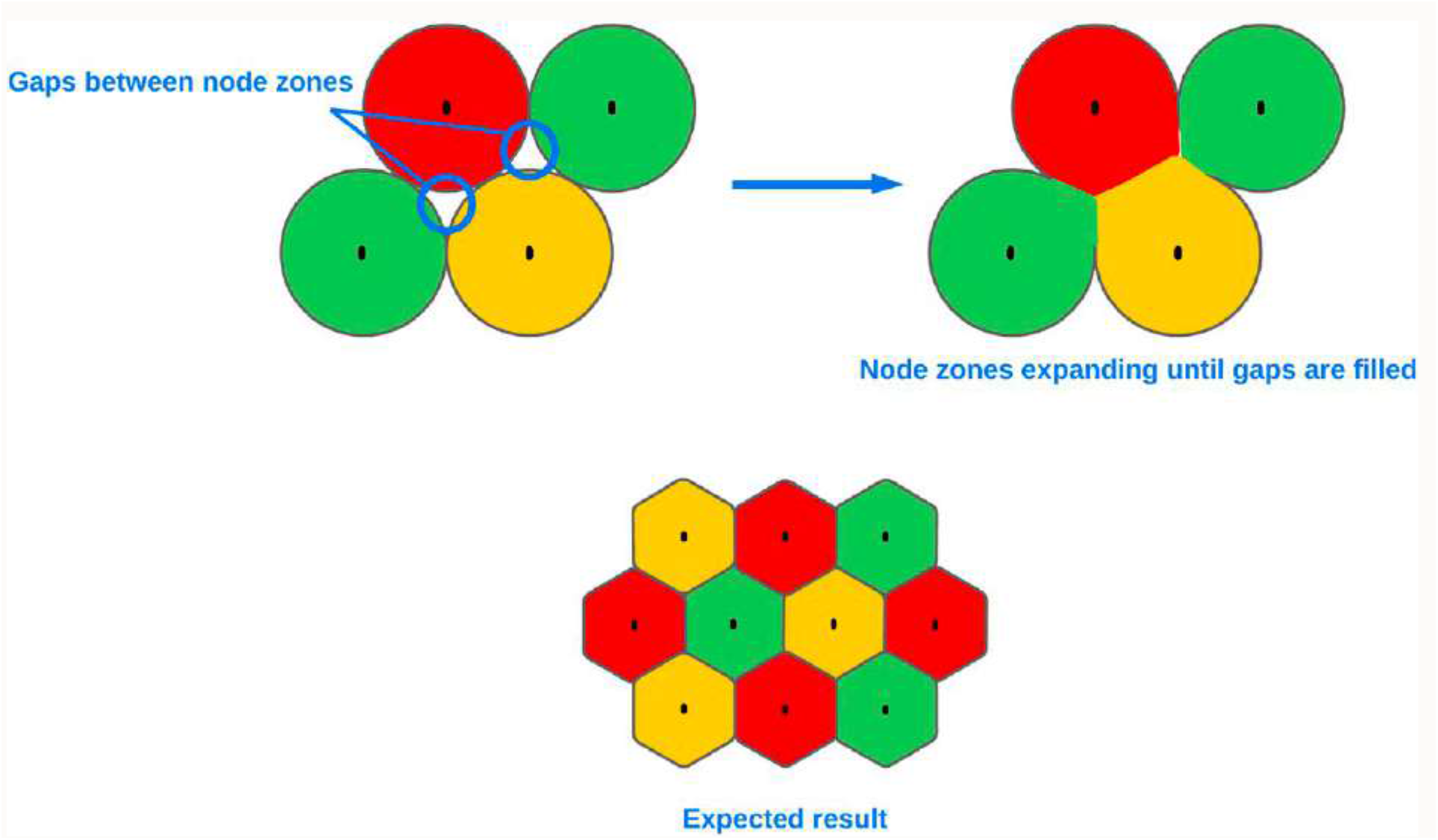
Expected emergent geometry from implementing a parallel Voronoi-like partition. Node zones spread outward and intersect uniformly, and each intersection between neighboring zones generates an edge between them. When a zone is surrounded and nodes of the same identity remain separated from one another, up to six intersections can occur around a given region, driving the emergence of approximately hexagonal domains. This behavior mirrors efficient hexagonal tilings observed in natural systems, such as packed water droplets or honeycomb structures^33^.

## 7. Future perspectives and conclusions

The proposed framework shows that a minimal combination of quorum-sensing-mediated signaling and CRISPR-Cas9-based genetic commitment can implement robust Voronoi-like spatial partitioning in a simulated bacterial colony. By assigning node identities through leader-follower dynamics and stabilizing these assignments via irreversible plasmid deletion, the system converts local signal exposure into persistent multicellular domains separated by well-defined interfaces^20,23^. Together with the quantitative clustering analysis, these results support the view that programmable spatial organization can be achieved in microbial populations using only local interactions, genetically encoded rules, and a small number of explicit spatial control parameters^6,16^.

Several limitations must be addressed for in vivo implementation. A primary constraint is the limited availability of orthogonal quorum-sensing systems and the risk of crosstalk between them, which complicates the use of fully independent signals for multiple node identities and increases metabolic burden^35^. Nevertheless, spatial signal reuse strategies based on sufficient separation between same-identity domains and sigmoidal input-output relationships may enable reuse of a given communication system across nonadjacent nodes^14,35^.

A second limitation is geometric: the current implementation and analysis are restricted to planar colonies, as the simulation environment operates in two dimensions^24,25^. Each simulated cell produces a defined output based on its local 2-dimensional environment, leading to a Voronoi-like partition in a plane, but it remains unclear how the same rules would behave in three dimensions. Extending the framework to 3-dimensional settings would require reconsidering diffusion, degradation, neighborhood definitions, and leader selection, and would likely reveal additional constraints and opportunities in tissues, biofilms, or structured biomaterials.

Overall, Voronoi-like spatial partitioning is positioned here as a representative problem for synthetic multicellular design rather than merely as a geometric construction. The leader–follower and CRISPR-based architecture demonstrates that distributed bacterial populations can be programmed to self-organize into stable, functionally differentiated domains using only local communication and irreversible commitment. Taken together with the explicit identification of key control parameters and robustness regimes, these results support the use of synthetic multicellular systems as minimal and controllable models for studying the core principles of spatial self-organization in microbial active matter^6,16,36^.

